# Thermal age, cytosine deamination and the veracity of 8,000 year old wheat DNA from sediments

**DOI:** 10.1101/032060

**Authors:** Logan Kistler, Oliver Smith, Roselyn Ware, Garry Momber, Richard Bates, Paul Garwood, Simon Fitch, Mark Pallen, Vincent Gaffney, Robin G. Allaby

## Abstract

Recently, the finding of 8,000 year old wheat DNA from submerged marine sediments (1) was challenged on the basis of a lack of signal of cytosine deamination relative to three other data sets generated from young samples of herbarium and museum specimens, and a 7,000 year old human skeleton preserved in a cave environment (2). The study used a new approach for low coverage data sets to which tools such as mapDamage cannot be applied to infer chemical damage patterns. Here we show from the analysis of 148 palaeogenomic data sets that the rate of cytosine deamination is a thermally correlated process, and that organellar generally shows higher rates of deamination than nuclear DNA in comparable environments. We categorize four clusters of deamination rates (α,β,γ,ε) that are associated with cold stable environments, cool but thermally fluctuating environments, and progressively warmer environments. These correlations show that the expected level of deamination in the sedaDNA would be extremely low. The low coverage approach to detect DNA damage by Weiss et al. (2) fails to identify damage samples from the cold class of deamination rates. Finally, different enzymes used in library preparation processes exhibit varying capability in reporting cytosine deamination damage in the 5’ region of fragments. The PCR enzyme used in the sedaDNA study would not have had the capability to report 5’ cytosine deamination, as they do not read over uracil residues, and signatures of damage would have better been sought at the 3’ end. The 8,000 year old sedaDNA matches both the thermal age prediction of fragmentation, and the expected level of cytosine deamination for the preservation environment. Given these facts and the use of rigorous controls these data meet the criteria of authentic ancient DNA to an extremely stringent level.

## Ancient DNA authentication by damage signal

Ancient DNA studies are providing an unprecedented level of insight into historical and prehistoric evolutionary processes (3). A critical aspect to these studies is the evaluation of the authenticity of ancient DNA. In the case of next generation sequencing data sets there is increased power to exploit signatures left by the processes of DNA decay (4). Principally, two decay characteristics are measured, the extent of DNA fragmentation and cytosine deamination.

It is thought that fragmentation is caused by hydrolysis of amino groups leading to loss of purine residues followed by strand cleavage by hydrolysis of the phosphate backbone. The rate of fragmentation has been strongly correlated with temperature over other possible influencing parameters giving rise to the concept of the thermal age of specimens (5,6). Consequently, temperature is the most informative predictor of DNA preservation and expected fragment length.

Cytosine deamination occurs prevalently towards the ends of fragments where there are single stranded overhangs and DNA bases are more exposed. The loss of amino groups results in the conversion of cytosine to uracil, and 5-methylcytosine to thymine. This leads to an elevation of C to T changes towards the end of fragments that can be measured in large data sets using applications such as mapDamage (7) and PMDtools (26). The processes that influence the rate of cytosine deamination are less well understood than in the case of DNA fragmentation, with a wide variation occurring between samples (3). Parameters that are known to be important include the length of the overhanging fragment (summarized as the λ value in mapDamage), and the propensity of cytosines and 5-methylcytosines in overhang regions to become modified to thymine, either directly or via a uracil template (summarized as the ∂s value in mapDamage). Generally, longer (and therefore thermally younger) DNA fragments are associated with larger overhangs that will contain more cytosine residues that are exposed.

## Cytosine deamination is a thermally correlated process

Currently, it is widely assumed that the rate of cytosine deamination most closely correlates to age and is largely resilient to the influence of temperature (8). On this basis there is an expectation of ancient DNA data to show greater signatures of cytosine deamination with age as a criterion of authenticity, although as of yet there has been no formal attempt to predict the level of cytosine deamination expected for a sample on the basis of age or other parameters.

Recent evidence has shown that the rate of deamination of 5-methylcytosine appears to be considerably higher at low latitudes associated with higher temperatures (9). Furthermore, previous correlations of cytosine deamination with age have not included permafrost samples that have shown very low ∂s values for their age (10). Such observations strongly suggest that the rate of cytosine deamination is influenced by thermal age rather than temporal age. We used mapDamage to re-analyse 72 human genomes obtained over a range of ages spanning several thousand years and latitudes spanning 39–59° (11), and a further 76 palaeogenomic data sets to see how well ∂s values correlate with age and temperature calculated from the WorldClim database, Figure 1. A clear correlation with temperature and age occurs within this group, with older samples from warmer environments showing a higher propensity for cytosine deamination (multiple r^2^ values 0.281 for nuclear DNA and 0.308 for organellar DNA). Notably, the correlation with temperature, but not age, is diminished if there are numerous samples from the same site under the same conditions, which is to be expected since the temperature parameter is nonvariant under these conditions. When corrected for this effect, the correlation with age and temperature becomes much stronger (median r value for nuclear 0.176, median r value for organellar 0.179) and the p value associated with temperature highly significant (5.08×10^−4^ for nuclear, and 3.92×10^−4^ for organellar DNA), Figure 1c. There is a stronger effect of temperature on deamination with organellar than nuclear DNA, which may reflect the lack of protection afforded to organellar DNA relative to nuclear DNA in the forms of chromatin structures.

**Figure 1.**
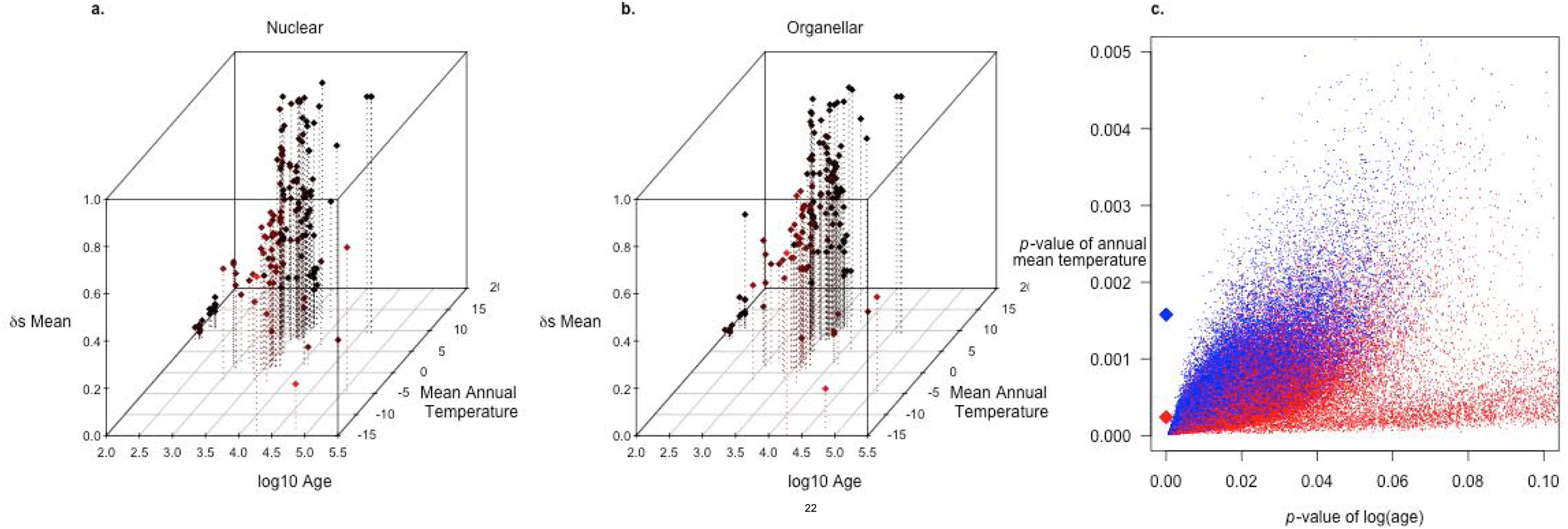
Trends of association of ∂S mean values from 148 palaeogenomic data sets (5) with temperature and age. A. Scatter plot of nuclear DNA data (association with age: p=9.4e-12; association with temperature temperature: p=0.0016; overall model p=4.24e-11; multiple r^2^ = 0.281. B. Scatter plot of organellar data (association with age: p=8.39e-13; temperature: p=0.0002; overall model p=2.50e-12; multiple r^2^ = 0.308). C. Distribution of p values for association with age and temperature for mean ∂S estimates from 100,000 random resamples (N=72) of 148 palaeogenomic data sets (5) to reduce the effect of multiple samples originating from the same site and therefore temperature conditions. Nuclear DNA shown in blue (association with age median p = 0.0152 [range 0.0028, 0.0495], association with temperature median p = 5.08e-4 [95% range 8.93e-5, 2.20e-3], multiple r^2^ = 0.176 [95% range 0.138, 0.221]). Organellar DNA shown in red (association with age median p = 0.0204 [95% range 0.0038, 0.1351], association with temperature median p = 3.92e-4 [95% range 7.27e-5, 1.79e-3], multiple r2 = 0.179 [95% range 0.139, 0.224]). Median values from panels A and B are plotted as diamonds.

We then calculated the rate of cytosine deamination required to result in the observed values of ∂s across 148 data sets from palaeogenomic studies, for nuclear and organellar DNA respectively, using an iterative algorithm to converge on a rate value to satisfy equation 1:

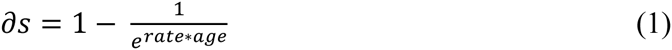

Strikingly, the observed cytosine deamination rates fall into a range over several orders of magnitude which we categorize into rate clusters (α,β,γ,ε), Figure 2a,b. The cluster with the lowest rates (a) has a mean rate of 1.08 × 10^−5^ and is made up of samples from cold environments (Table S1). The second group (β) which have a deamination rate an order of magnitude higher are made up of samples obtained from cool but variable environments in some cases, and sampled from petrous bones in others, with a mean of 6.49× 10^−5^. These samples include the Saqqaq Paleo-Eskimo (12), Columbian mammoths (13), Clovis-associated human from Montana (14), Ötzi, the Tyrolean “ice man” (15), and human samples from the Great Hungarian Plains (16). Most data sets fell within a larger cluster (γ) with rates an order of magnitude higher again and a mean rate of 1.9× 10^−4^ which encompasses the data sets we used in Figure 1. A fourth group of very high rates of cytosine deamination (ε) largely come from two studies (17,18) and are derived from plastid and mitochondrial DNA in high temperature environments, supporting the notion that organellar and plasmid DNA are more sensitive to the thermal component of cytosine deamination. In general, a very strong correlation is seen between rates of deamination and temperature of environment, Figure 2c,d. However, it is expected that variation rates will be modified positively with increasing amplitude of fluctuation in temperature diurnally, seasonally or over longer climate cycles. Furthermore, secondary issues of tissue types also have been observed to modify observed cytosine deamination rates (27,28).

**Figure 2.**
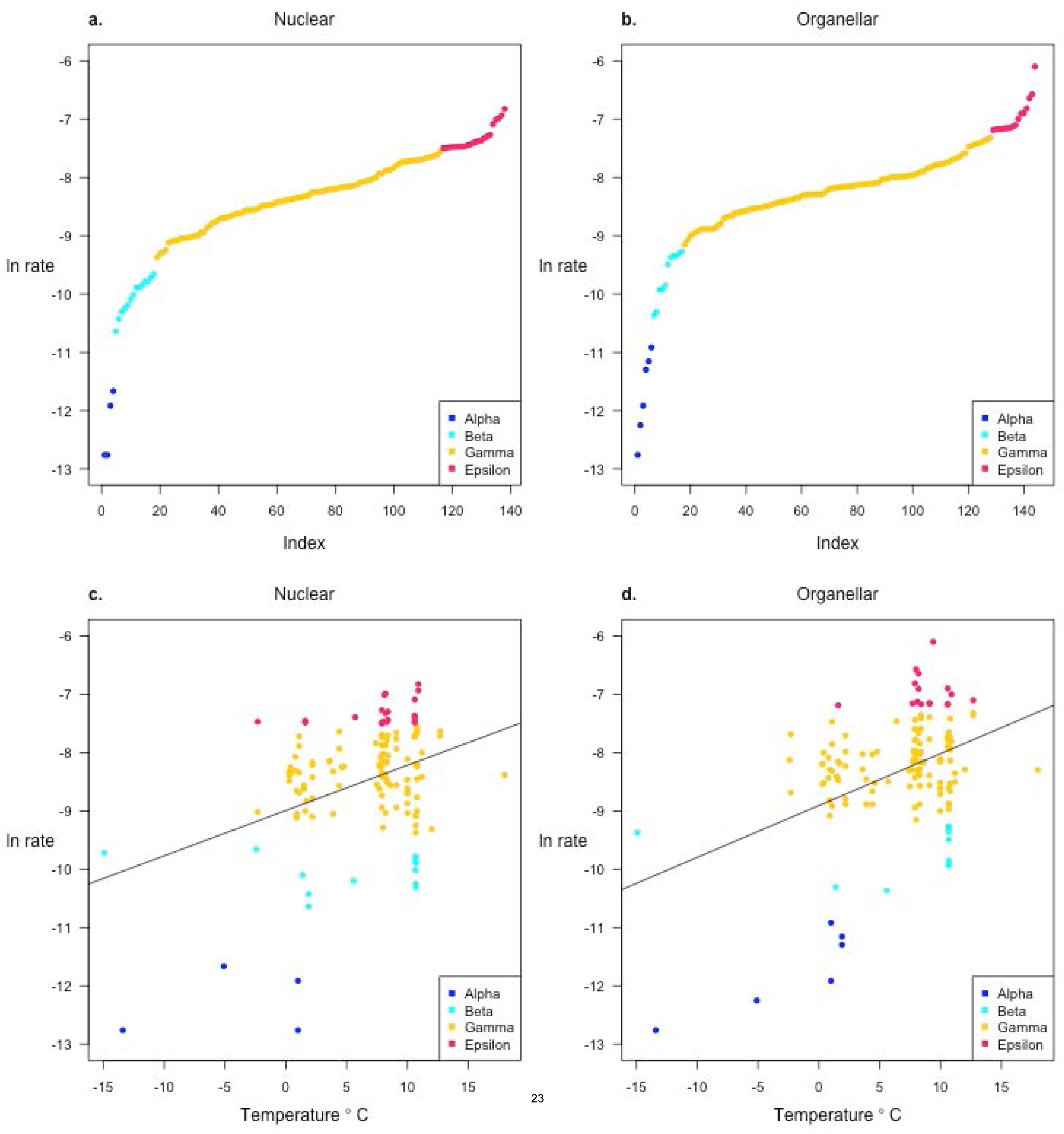
Cytosine deamination rates of 148 palaeogenomic studies. A. Natural logarithms of rates from nuclear DNA, B. rates from organellar DNA. Colours indicate the four rate classes, α,β,γ,ε in red, colour, colour and colour, respectively. C. Natural logarithms of nuclear rates against predicted temperature of environments, D. organellar rates against predicted temperature.

## Expected levels of cytosine deamination levels for different thermal classes of preservation

To date, there has been no formal approach published to establish expected cytosine deamination levels for given preservation conditions. To investigate what the expectation of the extent of cytosine deamination should be in a given temperature class of preservation context, we used the observed rates to calculate the mean and standard deviation for each of the fours classes of rate we identified. These were then used to estimate the range of ∂s values expected over time using equation 1, Figure 3. While the data show that even within a single class there is a finer scale of variation with temperature (Figure 1), these trends provide a convenient overall guide with which to assess the level of cytosine deamination that should be expected in samples. In the case of the sedaDNA sample, the preservation environment is a constant 4°C and has been so over the past 8,000 years. This is more constant than permafrost conditions, and certainly much more thermally stable than the environments in the samples of the higher beta category of deamination rates. In the rates we have measured, the most comparable sample to the sedaDNA environment we calculate to be a horse from the Taymyr peninsula (10), which is predicted to have a mean temperature of 1°C, and shows similar rates to permafrost samples within the alpha category of rates. The negative exponential nature of rate dependency on temperature apparent from Figure 2 leads to the expectation that cooler temperatures will vary less in deamination rates, it is therefore not surprising that the sedaDNA behaves in a way similar to permafrost samples. Consequently, the alpha group of rates is the likely most appropriate to apply for the sedaDNA, which leads to the expectation that after 8,000 years in this environment 4% of cytosines in exposed overhanging single strands of ancient DNA should have become deaminated, with a range of 2–6%, Figure 3. In the case of the 152 sedaDNA sequences that were identified as being wheat, and moderate 0.2 value for λ which leads to an average overhang of 3 bases, we would expect around 4.5 C to T events. In the sedaDNA data attributed to wheat there are 54 mismatches within 10 bases of the 3’ and 5’ termini. It therefore seems unlikely that this expected level of cytosine deamination would be detectable.

**Figure 3.**
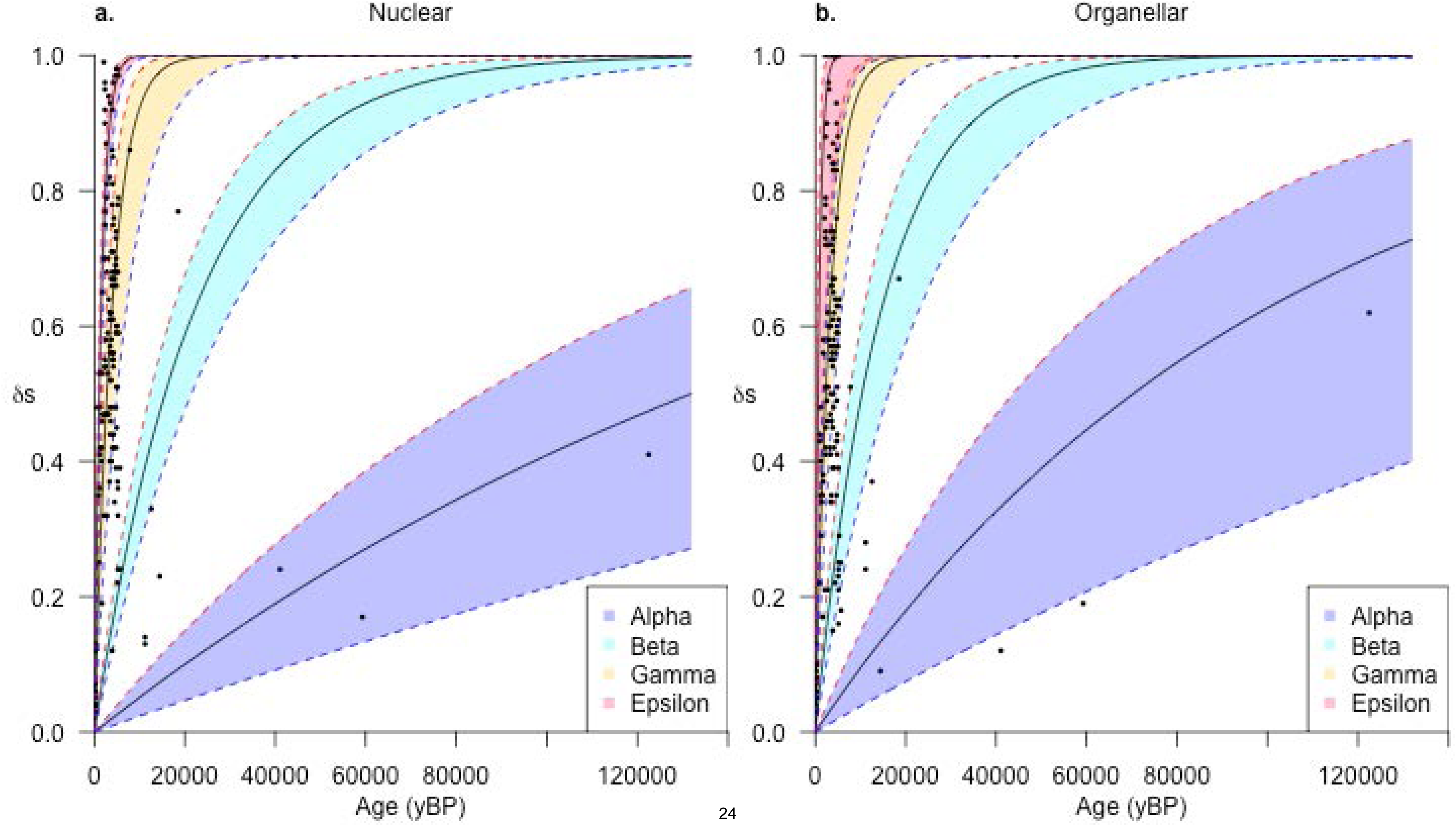
Cytosine deamination model for the prediction of ∂S values over time. Mean rates were calculated for each of the four rate classes a,b,g and e and used to plot ∂S over time using equation 1. Upper and lower bounds are described by one standard deviation from the mean. Observed ∂S values from 148 studies are plotted over the predicted trends. A. nuclear DNA predictions, B. organellar DNA predictions.

## Detectability of damage patterns in low coverage data across 148 data sets over multiple thermal classes

Weiß et al (2) demonstrated that their approach could not detect a signature of damage in the sedaDNA attributed to wheat from a single sampling of 152 sequences. The basis of the rejection of the wheat samples uses as criterion the level of goodness-of-fit to a negative exponential curve of cytosine-to-thymine mismatches from the edge of a DNA template, compared against a distribution of such values sampled in 152 read batches from potato herbarium samples. The wheat sedaDNA falls into the top 3% of values found in potato, giving a basis for rejection at a 0.05 significance level, for instance. This lack of detection of a damage signature matches our expectations for genuine ancient DNA of that age undergoing cytosine deamination in the alpha rate category. Notably, the test is highly dependent on the training set used to form the test distribution, about half of the test data sets of ‘known’ ancient DNA fail to produce distributions which place the sedaDNA in the 5% tail, despite its absence of damage signal (2).

Given the variance in deamination rates shown here, it is questionable whether an ancient DNA sample should be used as a test distribution for putative ancient DNA without respect to deamination rate comparability between the two. We explored how well this approach would work across the 148 data sets. Firstly, we sampled in 150-read batch sizes 10,000 times across each of the 148 data sets to calculate the goodness of fit *p* value to a negative exponential distribution for C to T transitions in the first 20 bases of a DNA fragment. From these data we produced empirical p distributions for all data sets, after the method described by Weiß (2). We then tested each sample against empirical distributions from all others, calculating whether 150-read batches would be rejected 5–10%, 10–20% or 20–100% of the time at the 0.05 level, Figure 4, Figure S1. We find that the potato and tomato *(Solanum* spp.) samples (19) used by Weiß et al. reject several known ancient DNA studies with a high probability, including the ~18,500 year-old M4 mammoth (13). We also find that samples from cool environments are rejected by many of the other studies, including the *Solanum* samples. Generally, many studies are rejected by many others with a high probability, and there is a tendency for warmer contexts to reject colder contexts even when the test samples from colder environments are much older, and generally for younger data sets to reject older ones. This suggests the test as applied here is highly prone to false negatives, and test sets should not be arbitrarily picked to validate damage patterns in other sample sets.

**Figure 4.**
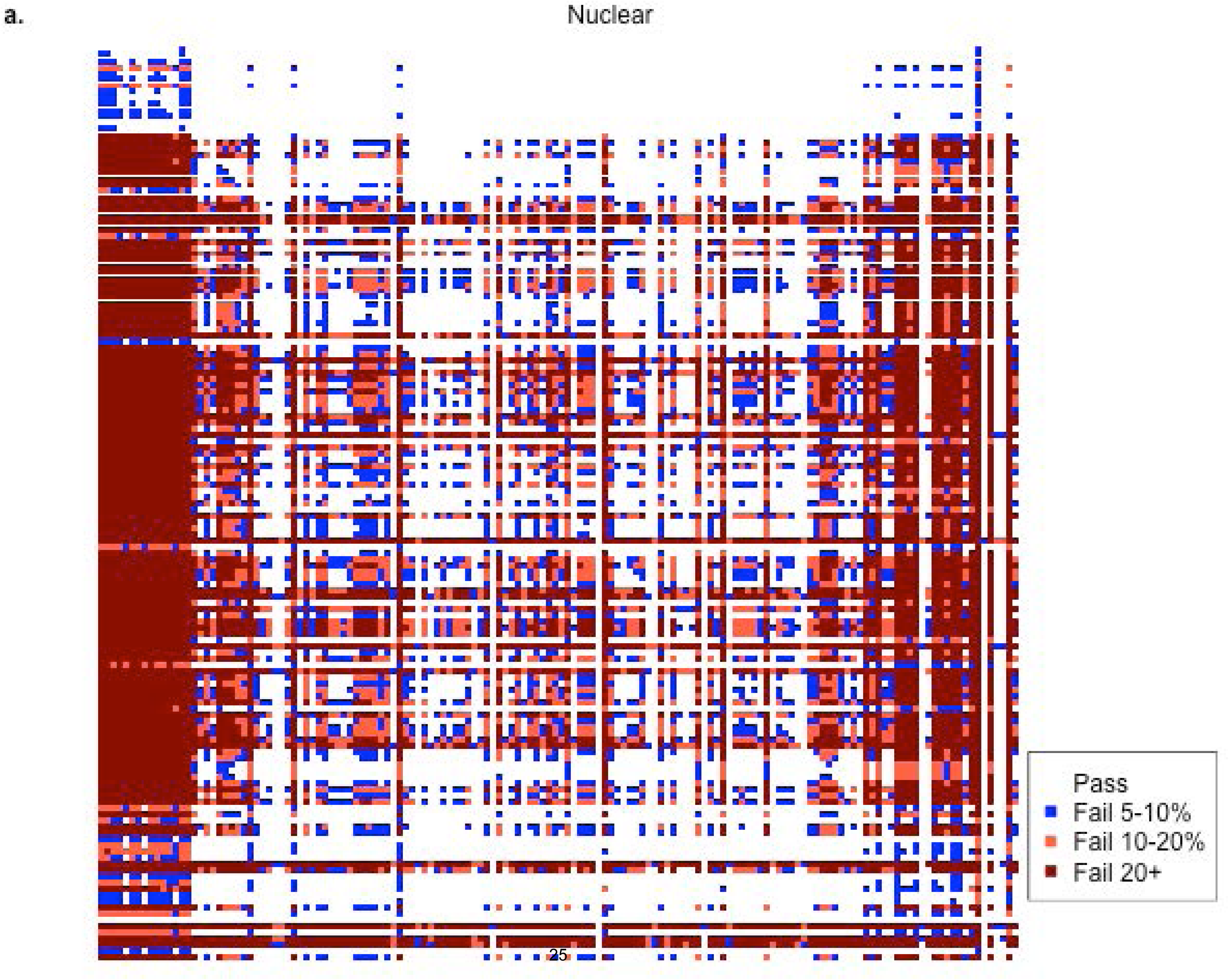
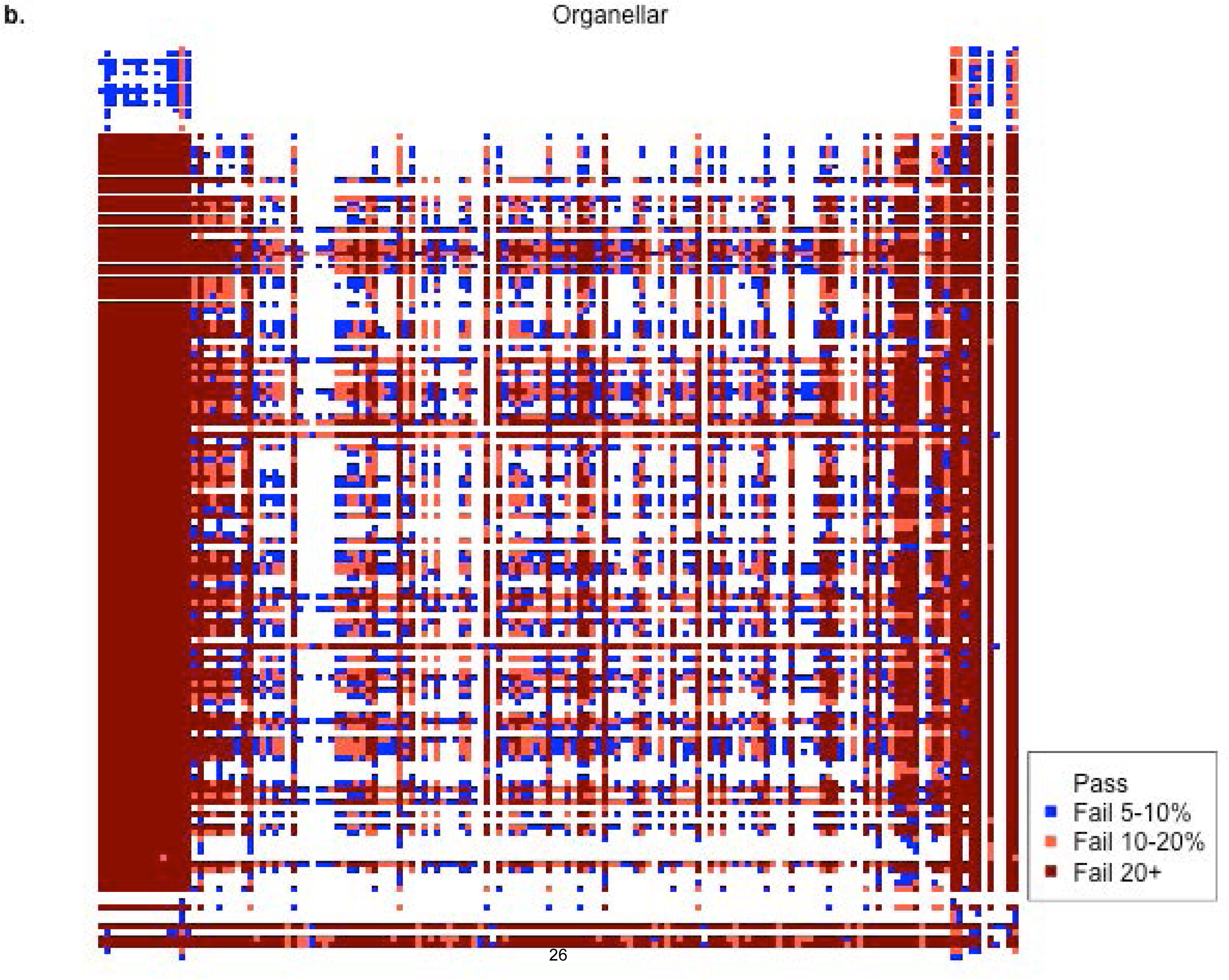
Cross-validation of 148 aDNA datasets using the method of Weiß et al (2). Empirical *p*-distributions generated from all samples (rows, age-sorted) were used to interrogate all other samples (columns, same order) to determine the percentage of 150-read subsamples that would fail aDNA validation at the 0.05 significance level. Colours indicate fail rates (rejection of ancient DNA authenticity), white, less than 5% chance of a batch being rejected, blue 5–10%, light red 10–20%, dark red 20–100%. Panel a nuclear DNA, panel b organellar DNA. E.g. a dark red cell indicates an instance where at least 20% of 10,000 subsamples generated from the column sample fail aDNA authentication against a *p*-distribution built from the row sample.

## Caveats in the detection of cytosine deamination rates

Complicating factors in the detection of cytosine deamination in ancient DNA are the protocols used to generate next generation sequence reads. A variety of polymerase enzymes are used, some of which have proofreading function and can replicate through uracil residues, some of which do not. Amplitaq Gold (company), Platinum *Taq* and *Pfx*,Hi Fidelity (life tech), and Accuprime *pfx* (company), all have proof reading activity, which means they are tolerant of uracils at the 5’ end of a DNA fragment. Conversely, *Pfu* does not have proof reading activity is intolerant to uracils at the 5’ end, and consequently damaged fragments of ancient DNA that contain uracils are not processed. In the case of the latter, library preparations based on this type of enzyme will not report a 5’ cytosine deamination profile if a directional library preparation method is utilized, such as Illumina’s y-tailed adapter system. The sedaDNA study was based on enzyme provided with the TruSeq Nano (Illumina) library preparation kit, and consequently the reads would not be expected to report a 5’ C to T damage profile. In the absence of a proof reading ability, damage profiles are still expected to be evident at the 3’ end as G to A mismatches. Consequently, in the case of the sedaDNA analysis it would have been more appropriate to test for an elevation of G to A mismatches at the 3’ end. However, given the alpha category of cytosine deamination rate, little or no signal is expected.

Even within the proofreading enzymes there is variability in the efficiency of damage reporting (20). Using data sets that were generated from the same samples using different protocols, we calculate that Amplitaq Gold reports a ∂s value 1.6 x higher than *Pfx*Accuprime *pfx* based on the La Braña genome study (21). Furthermore, different protocols will result in different ligation biases with sticky end ligations associated with bias against A’s in terminal regions (22).

## Discussion

Contrary to widely held view that the rate of cytosine deamination is a process only significantly correlated with age, analysis of the majority of shotgun palaeogenomic studies to date show a strong correlation also with temperature. The trends seen in these studies allow for some approximations to be made objectively of the expected extent of deamination to have occurred in any particular sample. The trends presented here may act as a guide in the application of criteria of authenticity, but also give some increased insight into the process of DNA diagenesis.

The sedaDNA from Bouldnor Cliff can be judged on two criteria of authenticity based on DNA decay processes, DNA fragmentation and cytosine deamination. In the case of DNA fragmentation, the pattern matches closely that expected using the thermal age tool (http://thermal-age.eu/) for a sample existing a constant temperature of 4°C for 8000 years. In fact, the sedaDNA appears to be slightly more degraded than expected with a median fragment length of about 115 bp as compared to the expected median of 134 bp. We find no difference in length between reads affiliated to wheat and reads assigned to other taxonomic groups in the palaeoenvironmental reconstruction suggesting that they have all been subject to the same fragmentation process. The expected extent of cytosine deamination for this preservation condition at the alpha rate is so low that detectability is not expected, and the pattern of C to T mismatches would not be expected to significantly follow a negative exponential pattern across a DNA fragment at this early stage of decay. These observations combined with the highly stringent controls applied in the sedaDNA study (1) provide compelling evidence of the authenticity of this ancient DNA.

## Methods

### Damage Estimation

We obtained unmapped (fastq) or mapped (bam) sequence reads from each of 148 publicly available ancient DNA datasets generated by shotgun sequencing without uracil removal by UDG, comprising anatomically modern humans (n=122), herbarium plant samples (n=16), Colombian and woolly mammoths (n=4), neandertals (n=3), horses (n=2), and polar bear (n=1). We avoided data generated through target capture experiments to avoid possible hybridization biases introduced by misincorporated residues. For unmapped samples, we used Flexbar (23) to trim adapter sequences in single-end read data, and PEAR (http://pear.php.net) to perform adapter trimming and read merging in paired-end datasets. We used the backtrack algorithm within the Burrowes-Wheeler Aligner (24) to align read data to the relevant reference genome (Table S1), and collapsed duplicates using the rmdup function in SAMtools (25). We filtered all bam files for a minimum mapping quality of 20 using the SAMtools “view” function, and filtered for minimum read length of 20 using Bash tools. We separated nuclear from organellar reads (mtDNA in mammals, plastid DNA in plants) and analyzed damage profiles independently. We used mapDamage 2.0 (7) to estimate damage parameters.

We used default settings for mapDamage in most cases, with the following exceptions: We downsampled large bam files to correspond with a 1 gigabyte input file (~10–20 million reads with typical dataset complexity and a human genome) using the mapDamage “-n” option. For libraries with much higher 3’ G-to-A mismatch than 5’ C-to-T mismatch indicating the use of a non-proofreading enzyme for library amplification (Table S1), we re-ran mapDamage with the “—reverse” option to estimate damage from the 3’ end only. We noted extremely high ∂s and λ values in the output from Mammoth M4 (13), which suggested a much higher rate of deamination than even much older permafrost samples. However, that library is dominated by very short fragments (Lynch et al 2015 supp), which we hypothesized could influence the mapDamage MCMC to over-estimate both ∂s and λ. We re-analyzed that sample considering only reads of ≥40nt, yielding the damage parameter values reported in Table S1. The Saqqaq data were mostly generated using a non-proofreading enzyme, but a small proportion of read files were reported to have been generated using a proofreading Platinum High Fidelity *Taq* polymerase (Life Technologies). We mapped all Saqqaq read files (n=218) to a human mitochondrial genome, used PMDtools (26) to generate misincorporation plots, and inspected these manually for elevated 5’ C-to-T mismatch, yielding two libraries apparently produced using a proofreading enzyme, one of which was used for analysis.

### Temperature Correlations

Temperature was estimated as follows: latitude and longitude were used to find annual mean temperature estimates for each of the samples. These were taken from the WorldClim current condition database (ref) using the R ‘raster’ package (version 2.4–18) at a resolution of 2.5 arc-minutes.

We used a linear model within the R statistical package to evaluate the relationship between DNA damage parameters, age, and temperature among all 148 samples, using the function: lm(∂s ~ log(age) + annual mean temperature).

### Age variance effect

For 50,000 iterations, we subsampled the dataset to include only one randomly-selected sample from any given pair of geographic coordinates, thus collapsing geographic sampling heterogeneity randomly to a single age value. We re-analyzed the ∂s–age–temperature effect in each iteration.

### Damage detection statistic from low coverage data

To test the sensitivity of the curve-fitting method described by Weiß *et al*. (2) to DNA degradation in a wide variety of preservation contexts, we generated 10000 random subsamples (sampled with replacement) of 150 reads from each of the 148 samples analyzed with mapDamage, and calculated mismatch profiles of the terminal 20nt using a custom perl script, after first downsampling very large files to approximately 10 million input reads using SAMtools (“-s” option in the “view” function). We considered only reads with full-length alignments and no indels. Mismatch summarizing was done in strand-aware fashion by direct comparison with the reference sequence, or by analysis of the MD:Z field as in PMDtools (26). We utilized the nls function in R to fit an exponential curve to terminal 5’ C-to-T or 3’ G-to-A mismatch (Table S1) according to the formula y ~ N*exp(-k*x), where *x* = base position, *y* = mismatch rate, and starting values for the rate parameter *k* and the modifier *N* were both invoked at 0.1. If the initial implementation failed to converge within the default limit of 50 iterations, we attempted up to 1000 additional combinations of starting *N* and *k* values sampled randomly from uniform distributions between 10e-10 and 10 (*N*) and -5 and 5 (*k*). Subsamples that fail to converge in all 1000 trials should most often be treated as failures to detect a significant exponential fit. However, conservatively, we excluded non-converging subsamples from any downstream analysis, and calculated goodness-of-fit *p*-value quantiles from only the set of converged subsamples.

## Acknowledgements

Thanks to Pontus Skoglund, Ludovic Orlando, Matthew Collins and Daniel Hebenstreit for comments. LK is supported by NERC Fellowship (NE/L012030/1), OS is supported by NERC (NE/L006847/1) and RW is supported by NERC. (NE/F000391/1).

